# Effects of Parkinson’s disease and dopamine on digit span measures of working memory

**DOI:** 10.1101/318311

**Authors:** John Patrick Grogan, Lisa Emily Knight, Laura Smith, Nerea Irigoras Izagirre, Alexandra Howat, Brogan Elizabeth Knight, Anastasia Bickerton, Hanna Kristiina Isotalus, Elizabeth Jane Coulthard

## Abstract

**Rationale:** Parkinson’s disease (PD) impairs working memory (WM) - the ability to maintain items in memory for short periods of time and manipulate them. There is conflicting evidence on the nature of the deficits caused by the disease, and the potential beneficial and detrimental effects of dopaminergic medication on different WM processes.

**Objectives:** We hypothesised that PD impairs both maintenance and manipulation of items in WM and dopaminergic medications improve this in PD patients but impair it in healthy older adults.

**Methods:** We tested 68 PD patients ON and OFF their dopaminergic medication, 83 healthy age-matched controls, and 30 healthy older adults after placebo and levodopa administration. We used the digit span, a WM test with three components (forwards, backwards and sequence recall) that differ in the amount of manipulation required. We analysed the maximum spans and the percentage of lists correctly recalled, which probe capacity of WM and the accuracy of the memory processes within this capacity, respectively.

**Results:** PD patients had lower WM capacity across all three digit span components, but only showed reduced percentage accuracy on the components requiring manipulation (backwards and sequence spans). Dopaminergic medication did not affect performance in PD patients. In healthy older adults, levodopa did not affect capacity, but did impair accuracy on one of the manipulation components (sequence), without affecting the other (backwards).

**Conclusions:** This suggests a non-dopaminergic deficit of maintenance capacity and manipulation accuracy in PD patients, and a potential “overdosing” of intact manipulation mechanisms in healthy older adults by levodopa.

## INTRODUCTION

Working memory (WM) involves the maintenance and manipulation of elements held in memory for short periods of time. Early models suggested WM is composed of phonological and visuospatial storage components, and a central executive that manages attentional demands and the manipulation of stored elements (Baddeley, 2012).

Parkinson’s disease (PD) patients have impaired WM, especially for visuospatial tasks or complex tasks requiring manipulation (Lewis, Slabosz, Robbins, Barker, & Owen, 2005). Maintenance of WM is also impaired in PD (Fallon, Mattiesing, Muhammed, Manohar, & Husain, 2017), though a meta-analysis of 56 WM span studies suggested that verbal maintenance was reduced to a lesser extent than verbal manipulation, and that spatial WM was impaired the most (Siegert, Weatherall, Taylor, & Abernethy, 2008). Dopaminergic medication might improve both maintenance and manipulation in PD patients (Beato et al., 2008; Owen, Iddon, Hodges, Summers, & Robbins, 1997; Zokaei, Burnett Heyes, Gorgoraptis, Budhdeo, & Husain, 2015), though some tasks have found no effect of medication (Fournet, Moreaud, Roulin, Naegele, & Pellat, 2000; Zokaei et al., 2015) or only shown benefits in specific subgroups such as patients with low baseline WM (Warden, Hwang, Marshall, Fenesy, & Poston, 2016) or in patients with onset of motor symptoms on their left-side (Hanna-Pladdy, Pahwa, & Lyons, 2015). Importantly, many of these studies may have been underpowered, with small sample sizes (n=7-28) exacerbated by between-subjects designs, which may contribute to the conflicting results.

Dopamine can also impair WM, perhaps due to an overdosing of intact areas of the brain (Cools & D’Esposito, 2011). This is more often seen in healthy participants than patients (Bloemendaal et al., 2014; Fallon, van der Schaaf, Ter Huurne, & Cools, 2016), although benefits of dopamine are also seen (Fallon et al., 2016; Naef et al., 2017) as are no-effects (Linssen, Vuurman, Sambeth, & Riedel, 2012); this may reflect different optimal levels of dopamine for different functions within WM such as maintenance and manipulation. Interestingly, in one of these studies testing WM following administration of methylphenidate (a dopamine and noradrenaline reuptake inhibitor), both beneficial and deleterious effects were demonstrated in the same group of participants (Fallon et al., 2016); methylphenidate improved distractor resistance on a spatial WM task, but impaired flexible updating of information held in WM. This suggests that the ‘overdose’ effects reported in some studies may not be seen when using broad measures of WM but require specific measurement of the different underlying processes.

We hypothesised that PD would impair maintenance and manipulation of WM, and that dopaminergic medication would remediate the deficits in PD patients while ‘overdosing’ and impairing WM in healthy older adults. We used a simple WM measure, the digit span (Wechsler, 2008), which is commonly given in neuropsychological assessments. We used three variations of the digit span (forwards, backwards, sequence recall) with different contributions of maintenance and manipulation processes. This was given to a large sample of PD patients ON and OFF their normal dopaminergic medication and healthy age-matched controls (HC), and a separate group of healthy older adults after administration of placebo or levodopa.

## METHODS

### Ethical approval

Data are presented from several different studies running under different ethical approvals. Experiment 1 studies were approved by Frenchay and Southwest Central Bristol NHS RECs. Experiment 2 was approved by University of Bristol Faculty REC. All procedures were carried out in accordance with the relevant guidelines and regulations. All participants gave written informed consent, in accordance with the Declaration of Helsinki.

### Participants

#### Experiment 1

Demographic details for all groups are presented in Table 1.

**Table 1.** Demographics of participants tested in experiments 1 and 2. Statistical comparisons are against HC from Experiment 1 (standard deviations in parentheses). The older adults from Experiment 2 did not differ from the HC in Experiment 1 on any measure (p>.05). *p < .05 for PD vs HC, ^+^p < .05 for PD ON vs PD OFF. MoCA= Montreal Cognitive Assessment, DASS=Depression, Anxiety and Stress Scale, BIS=Barratt Impulsivity Scale, LARS=Lille Apathy Rating Scale, LDE=Levodopa Dose Equivalency, MDS-UPDRS-III=Movement Disorder Society Unified Parkinson’s Disease Rating Scale section III.

| Measure | PD Experiment 1 | HC Experiment 1 | HC Experiment 2 |
| --- | --- | --- | --- |
| Number | 68 | 83 | 30 |
| Age | 68.40 (6.53) | 69.66 (8.49) | 70.67 (6.83) |
| Gender (M:F) | 49:19* | 39:44 | 14:16 |
| MoCA | 25.31 (2.88)* | 26.94 (2.37) | 26.23 (3.15) |
| Years Education | 13.70 (3.19) | 14.65 (2.87) | 14.33 (3.48) |
| DASS | 22.91 (16.78)* | 14.07 (18.27) | 11.27 (10.29) |
| BIS | 54.04 (8.50) | 50.33 (9.00) | 48.50 (8.54) |
| LARS | -22.71 (4.81)* | -27.28 (4.57) | -26.60 (5.54) |
| LDE | 605.41 (353.89) |  |  |
| Years Symptoms | 5.79 (3.86) |  |  |
| Years Diagnosed | 4.63 (3.47) |  |  |
| MDS-UPDRS-III ON | 27.95 (13.39)<sup>+</sup> |  |  |
| MDS-UPDRS-III OFF | 32.87 (13.41) |  |  |

To generate a good sample size, we applied a robust, consistent protocol for testing digit span across several different studies. In total, we collected data from 68 PD patients and 83 HC who all performed the digit span (as well as other cognitive tasks dependent on study). Patients with a diagnosis of idiopathic PD were recruited from neurology and movement disorder clinics at Southmead and Frenchay Hospitals in Bristol, UK. All were taking levodopa and/or dopamine receptor agonists, were not taking irreversible mono-amine oxidase inhibitors or acetylcholinesterase inhibitors, and did not have deep-brain stimulators implanted. They had no serious neurological disorders other than PD and had normal or corrected vision and hearing.

HC were recruited from our healthy volunteer database. They were 55 years or older, had no neurological disorders, were not taking dopaminergic medications, and had normal or corrected to normal vision and hearing.

#### Experiment 2

We recruited 35 healthy older (65+ years) adults from volunteer databases and Join Dementia Research databases. The same inclusion/exclusion criteria as for the HC group above were used, as well as contraindications and medical exclusions pertaining to the drugs administered (see Supplementary Materials 1 for full exclusion criteria). Two participants withdrew before completing one session, and three withdrew before completing both the drug and placebo session, leaving data from 30 participants analysed here.

### Procedure

In the digit span, the experimenter reads aloud a list of digits at a rate of 1 per second and the participant must repeat the list back. All digits must be in the correct order for the list to be marked correct. The lists start at a length of 2 digits, and two lists of each length are read out. The list lengths increase by 1 digit until the participant gets both lists of the same length correct.

There are three components of the digit span: in the *forwards* span, the list must be recalled in the same order as said by the experimenter; in the *backwards* span, participants must repeat it in the reverse order to presentation order; in the *sequence* span, participants must recall the list in ascending numerical order. Forwards and sequence components present two lists of each length from 2-9 digits. The backwards component presents four lists of 2 digits length, then two lists from lengths of 3-8 digits. There are 16 lists in total for each component.

#### Experiment 1

PD patients completed the three components (forwards, backwards, sequence) of the digit span once ON and once OFF medication (medication order randomised and counterbalanced), while HC completed it once. When coming OFF medication, PD patients were withdrawn from standard release dopaminergic medication for a minimum of 16 hours and from long-lasting dopaminergic medications for a minimum of 24 hours.

#### Experiment 2

This was a within-subjects double-blinded, placebo-controlled study. The participants completed both the drug and placebo conditions, in a randomised, counterbalanced order.

Healthy older adults received 10mg/ml domperidone or a placebo, followed by 187.5mg co-beneldopa (150mg levodopa, 37.5mg benserazide) or a second placebo. Neither participant nor experimenter knew on which visit the participant received the drug or placebo. Their heart rate and blood pressure were monitored before and after administration. After 1.5 hours they completed the digit span, along with other cognitive tests.

### Data analysis

We used several scoring measures for the digit span to capture different sources of errors. The maximum span length correctly recalled gives a measure of the maximum capacity of a participant’s WM. This is calculated for each span component (forwards, backwards and sequence) separately.

People can also make errors even before they have hit their capacity limit, which is not picked up by the maximum span measure. There are several measures sensitive to the number of errors in WM which reflect WM *accuracy* rather than *capacity*. The number of lists recalled correctly gives a simple count of these errors but is confounded by the fact that people with smaller capacities will exit the test earlier and thus not attempt as many lists as someone with a larger span. Therefore, we analysed the percentage of lists recalled correctly, which corrects for the number of lists attempted and gives a more reliable and accurate measure of the accuracy of WM (Conway et al., 2005; Friedman & Miyake, 2005).

Assessing the percentage of digits recalled correctly rather than the lists may have even greater reliability and sensitivity as it captures extra information in the data (Friedman & Miyake, 2005; Unsworth & Engle, 2007). Unfortunately, the exact digits recalled were not consistently recorded for all participants; there are only digit error data for 45 PD and 52 HC from Experiment 1, and only for two participants from Experiment 2. These data are presented in Supplementary Materials 2 but are not reported here due to the much lower power and weaker effects.

Q-Q plots showed the spans were all approximately normal. Between-subject ANOVAs and t-tests were used to compare PD patients and HC, and within-subject t-tests to compare the effect of medications on PD patients and the effect of levodopa on healthy participants in Experiment 2. When comparing the three components, Bonferroni corrections were applied (α=0.0167). Data were analysed using SPSS (IBM, version 23.0).

### Data availability

Experiment 1 did not obtain consent to share individual participants’ data, so we are not able to publish or provide the data without further permission from our study sponsor.

Anonymised data from Experiment 2 are available from the University of Bristol’s data repository, data.bris, at https://doi.org/10.5523/bris.15du56inneqal1ys8rhzuhbmmu.

## RESULTS

### Experiment 1

PD patients had lower maximum spans for forwards (F (2, 216) = 4.572, p = .011, *η*^2^_*p*_ = .041), backwards (F (2, 216) = 6.590, p = .002, *η*^2^_*p*_ = .058), and sequence (F (2, 216) = 8.317, p < .001, *η*^2^_*p*_ = .072) spans, with greater effect sizes in the manipulation spans (backwards and sequence spans; see Fig 1 and Table 2). Paired t-tests showed no significant differences between PD ON and OFF dopaminergic medication on any span (forwards: t (67) = 0.944, p = .348, *d* = 0.102; backwards: t (67) = -.309, p = .758, *d* = 0.032; sequence: t (67) = 0.456, p = .650, *d* = 0.049).

**Fig 1.**
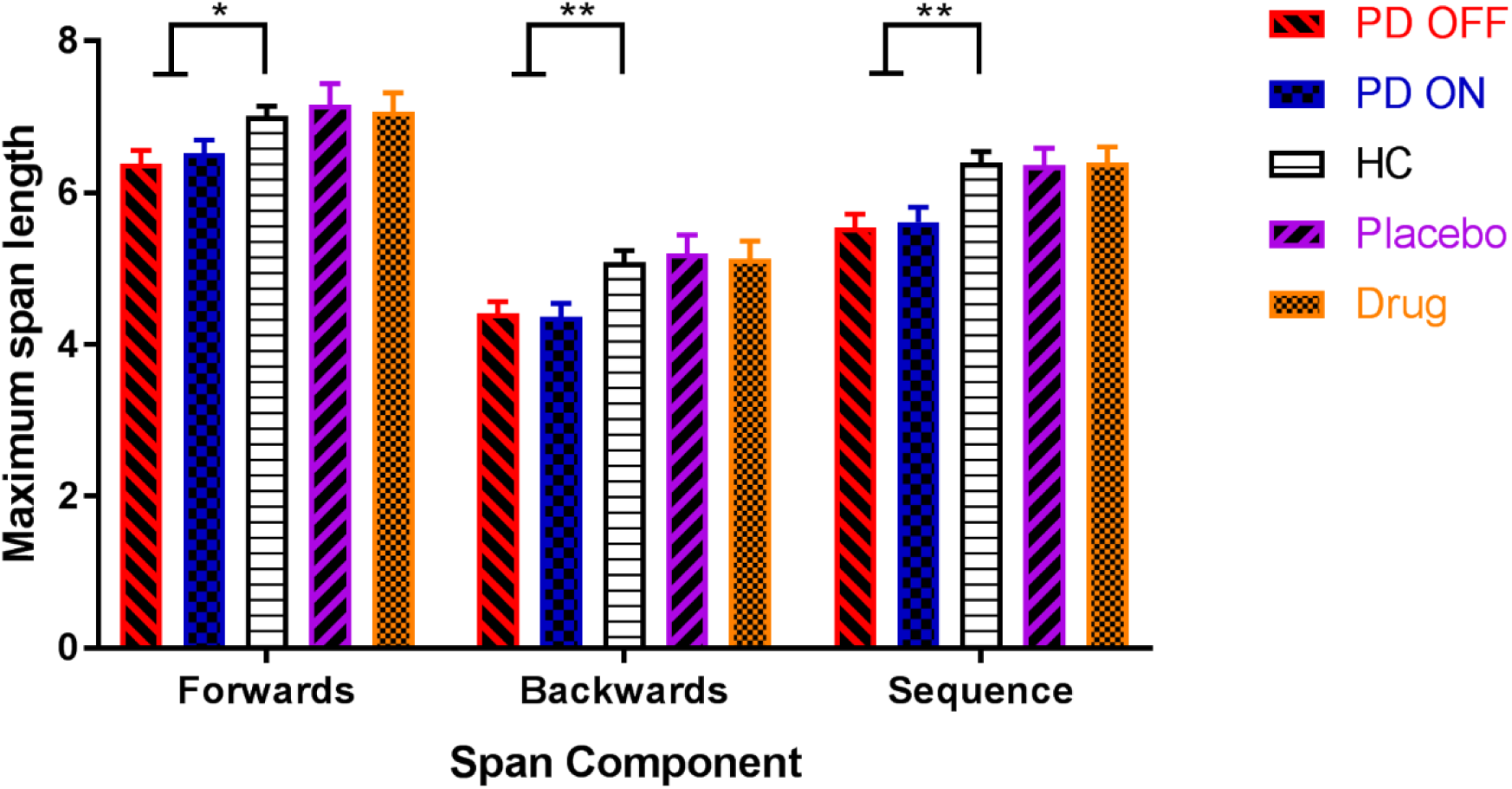
The mean maximum spans for PD patients ON and OFF dopamine, HC, and healthy older participants on levodopa and placebo on each component of the digit span (SEM bars). PD patients had lower maximum spans than HC for all span components, but there were no effects of dopamine for PD patients or healthy older adults for any component. *p < .01667 (Bonferroni-corrected threshold), **p < .001667.

**Table 2.** Effect sizes and p-values from group comparisons of digit span measures. Between-subject one-way ANOVAs were used to compare HC vs PD ON vs PD OFF, while paired t-tests were used to compare PD ON vs PD OFF and Drug vs Placebo. Bonferroni corrections were applied at a significance threshold of α=.01667. *p < .0167, **p < .00167, ***p < .000167.

| Comparison | Measure | Forwards |  | Backwards |  | Sequence |  |
| --- | --- | --- | --- | --- | --- | --- | --- |
|  |  | d | p value | d | p value | d | p value |
| **HC vs PD-ON vs PD-OFF** | Max Span | 0.414 | 0.011358* | 0.496 | 0.001667** | 0.557 | 0.000331** |
|  | % lists correct | 0.263 | 0.164839 | 0.578 | 0.000167*** | 0.487 | 0.001975* |
| **PD-ON vs PD-OFF** | Max Span | 0.102 | 0.348482 | 0.032 | 0.758273 | 0.049 | 0.650100 |
|  | % lists correct | 0.147 | 0.220807 | 0.281 | 0.048302 | 0.167 | 0.155641 |
| **Drug vs Placebo** | Max Span | 0.070 | 0.609901 | 0.051 | 0.757700 | 0.028 | 0.902247 |
|  | % lists correct | 0.072 | 0.654236 | 0.085 | 0.654438 | 0.689 | 0.006718* |

PD patients made more errors (i.e. lower percentage of lists correct) than HC only for the manipulation spans (see Fig 2; backwards: F (2, 216) = 9.060, p = .0002, *η*^2^_*p*_ = .077; sequence: F (2, 216) = 6.410, p = .0020, *η*^2^_*p*_ = .056) but not for the forwards span (F (2,216) = 1.818, p = .1648, *η*^2^_*p*_ = .017). Dopaminergic medication did not affect the percentage of lists correct (forwards: t (67) = -1.236, p = .2208, *d* = 0.147; backwards: t (67) = 2.011, p = .0483, *d* = 0.281; sequence: t (67) = 1.436, p = .1556, *d* = 0.167).

**Fig 2.**
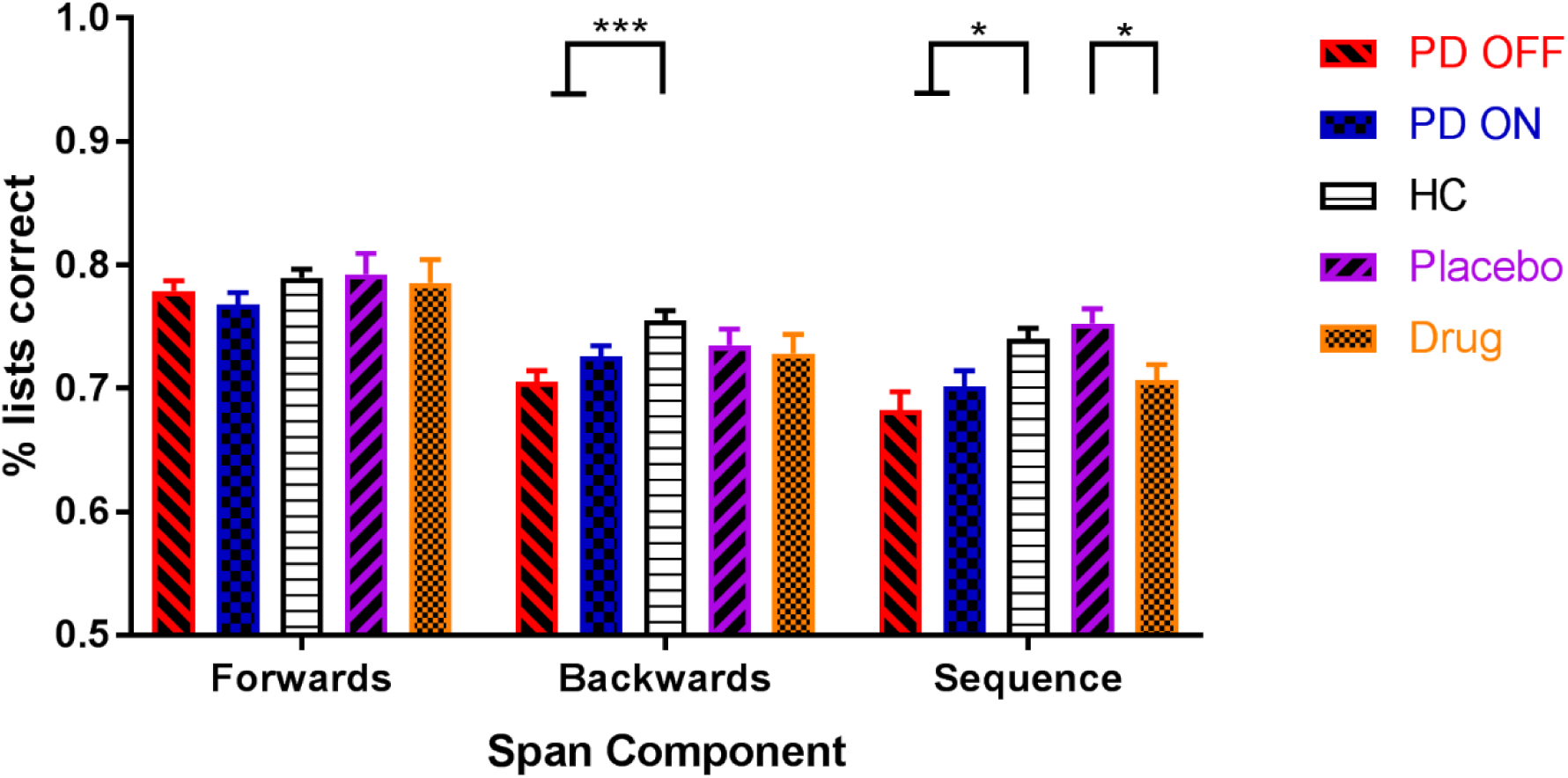
The mean percentage of lists correct for each group, on each digit span component (SEM bars). PD patients had lower accuracy scores for the backwards and sequence components but not the forwards component. Dopamine did not affect accuracy in PD patients, but levodopa did decrease accuracy on the sequence component for healthy older adults. *p < .01667 (Bonferroni-corrected threshold), ***p < .0001667.

As PD patients had lower maximum spans on the forwards component, but did not have lower percentage of lists correct, we compared these two measures directly. We converted maximum span length into a percentage to be on the same scale as the percentage of lists correct and ran a mixed ANOVA (within-subject factor: measure type, between-subject factor: group (PD or HC)). The forwards component had a significant measure*group interaction (F (1, 217) = 6.511, p = .011, *η*^2^_*p*_ = .029) that passed the Bonferroni-corrected threshold (α = .01667), while backwards (F (1, 217) = 4.641, p = .032, *η*^2^_*p*_ = .021) and sequence (F (1, 217) = 4.984, p = .027, *η*^2^_*p*_ = .022) did not. This suggests that PD affects the maximum span length and percentage of lists correct differently for the forwards component, but not the backwards or sequence components.

No effects of levodopa dose equivalency were seen, nor were there effects of laterality of motor symptoms or medication type (see Supplementary Materials 2).

### Experiment 2

Looking at the healthy older adults given levodopa and placebo, levodopa did not affect maximum span digit span component (forwards: t (29) = 0.516, p = .610, *d* = 0.070; backwards: t (29) = 0.311, p = .758, *d* = .051; sequence: t (29) = -.0124, p = .920, *d* = 0.028) (see Fig 1 and Table 2).

However, levodopa did decrease the percentage of lists correctly recalled for the sequence component (t (29) = -2.919, p = .007, *d* = 0.689), though not for the forwards (t (29) = -0.453, p = .654, *d* = 0.072) or backwards (t (29) = -0.452, p = .654, *d* = 0.085) components (see Fig 2). No effects of weight-adjusted levodopa dose were seen (see Supplementary Materials 2).

Only two participants from Experiment 2 had their error responses recorded, so the percentage of digits correct were not analysed for Experiment 2.

## DISCUSSION

PD patients had lower maximum spans for each component of the digit span, as well as lower percentage of lists recalled correctly for backwards and sequence spans.

Dopaminergic medication did not improve performance on any component or measure in PD patients. In healthy older adults, levodopa did not affect maximum span, but did decrease the percentage of lists correctly recalled for the sequence span.

PD patients were only impaired on the maintenance component (forwards digit span) when measuring the maximum span length recalled, not the percentage of lists correct. However, for the manipulation components (backwards and sequence spans) PD impaired the maximum capacity and percentage of lists correct similarly. This distinction suggests that the two measures are tapping into distinct processes during WM – the capacity and the accuracy. It also suggests that PD impairs the capacity for maintenance and manipulation processes, but only reduces accuracy of manipulation processes. This aligns with previous literature which has suggested that while PD does lead to more decay of precision of items held in memory (Fallon et al., 2017; Zokaei et al., 2015), there are greater deficits when manipulation is required (Lewis et al., 2005) and that this is due to increased number of errors (Fallon et al., 2017).

Alternatively, the difference in the measures may simply reflect poorer sensitivity of the percentage list accuracy measure. However, previous literature suggests the percentage accuracy scores actually have *greater* sensitivity and reliability than simpler measures such as the maximum span length (Conway et al., 2005; Friedman & Miyake, 2005), which would argue against this view. Additionally, PD OFF scored (non-significantly) higher on the percentage of lists measure than PD ON, a different direction of effect to the maximum span measure. This suggests that the lack of effect is not simply due to smaller differences between the groups but may reflect different processes underlying the data.

The general deficit in WM capacity could reflect a reduction in the number of items that can be maintained in WM having a knock-on effect onto the manipulation components in the backwards and sequence spans. If PD reduces the number of items a person can hold in their memory, then this would also reduce the maximum spans possible in the backwards and sequence components. This is unlikely to be the sole driver of this deficit however, as backwards and sequence components had lower mean maximum spans than the forwards component, meaning they were not hitting the ceiling imposed by the maintenance capacity and that there is an extra source of error in these manipulation spans.

If PD harms WM but dopaminergic medication does not improve it, then a non-dopaminergic pathology is suspected. PD patients have alterations to many neuromodulatory systems including noradrenaline, acetylcholine and serotonin (Jellinger, 1991; Scatton, Javoy-Agid, Rouquier, Dubois, & Agid, 1983), which may underlie the deficit. Alternatively, it could be a dopaminergic pathology, but simply one that is too severe to be repaired by dopamine replacement therapy, although this seems unlikely given that motor symptoms, usually seen before cognitive changes, are still helped by dopaminergic medication, as evident in the reduced UPDRS scores in PD patients when ON medication.

More interesting is the apparent sparing of maintenance accuracy from the WM deficit caused by PD. This could suggest that the underlying processes accounting for errors on the spans is different when manipulation of the items is required, as PD seems to reduce the maximum number of items that can be maintained in WM, without increasing errors made. To explain this, we invoke the multicompartment model of WM (Baddeley, 2003), which posits a phonological loop for storage of items, and a central executive that mediates manipulation of items stored. We propose that PD impairs the capacity of the phonological loop, without increasing errors in storage under this limit, and also impairs the central executive’s ability to interact with items stored.

The pattern of results from Experiment 2 suggest that levodopa does not affect the maximum capacity of any of the span types but may reduce the percentage of lists recalled correctly only for the sequence span. This induced deficit supports the dopamine overdose hypothesis which posits that dopaminergic drugs will overdose intact functioning in the brain, leading to impairments (Cools & D’Esposito, 2011). That manipulation accuracy was reduced by levodopa corroborates reports that methylphenidate impairs the flexible updating of WM information (Fallon et al., 2016), which would be needed in the sequence span. However, this effect should be interpreted with caution. Unlike the pattern of effects seen in PD patients, this one is isolated. There were no impairments on the other manipulation component (the backwards span); indeed, the percentage of lists correct for backwards span was not even close to significantly different. Therefore, while it may be that levodopa does impair manipulation accuracy of WM items, it is also possible that this is simply an artefact or false positive.

There are several drawbacks in using the digit span that should be considered. As only two lists of each length are presented, it provides a very noisy measure of performance. Adapted versions presented via computer are available which use give repeated presentations of list lengths, and do not quit when they fail to recall the lists but instead decrease the length and then increase it back up if they recall that one correctly (Woods et al., 2011). This step-up/step-down procedure is more sensitive to people’s maximum capacities. Computerised assessment would also rule out any variability induced by slightly different speaking speeds, accents, volume, and diction, from different experimenters, which may have affected performance. Other computerised tasks are available which provide analogue error measures on WM, which have shown far greater sensitivity than the digit span (Zokaei et al., 2015). Work with these tasks has suggested that WM capacity may not be determined by the number of discrete ‘slots’ for information but rather by the allocation of a shared capacity resource (Bays, Catalao, & Husain, 2009; Schneegans & Bays, 2016). Future work with these more sensitive tasks will be able to separate out the specific WM processes impaired by PD and affected by dopamine.

In summary, PD impaired maximum capacity of maintenance spans, and the capacity and accuracy of manipulation spans. Despite this, PD patients show no benefit of dopaminergic medication on maintenance or manipulation of WM, suggesting a non-dopaminergic deficiency. Levodopa also did not affect WM capacity in healthy older adults, but may have decreased accuracy on manipulation span, although this effect should be viewed with caution.

## ACKNOWLEDGEMENTS

The authors thank BRACE for the use of their building, all the patients and participants who took part in this research, and the funders (Wellcome [097081/Z/11/Z] and Bristol Research into Alzheimer’s and Care of the Elderly charity [BRI17/15]).

